# Chaperone AMPylation modulates aggregation and toxicity of neurodegenerative disease-associated polypeptides

**DOI:** 10.1101/183723

**Authors:** Matthias C. Truttmann, David Pincus, Hidde L. Ploegh

## Abstract

Proteostasis is critical to maintain organismal viability, a process counteracted by aging-dependent protein aggregation. Chaperones of the heat shock protein (HSP) family help control proteostasis by reducing the burden of unfolded proteins. They also oversee the formation of protein aggregates. Here, we explore how AMPylation – a post-translational protein modification that has emerged as a powerful modulator of HSP70 activity – influences the dynamics of protein aggregation. We find that adjustments of cellular AMPylation levels in *C.elegans* directly affect aggregation properties and associated toxicity of amyloid-β (Aβ), of a polyglutamine (polyQ)- extended polypeptide and of α-synuclein (α-syn). Expression of a constitutively active *C. elegans* AMPylase Fic-1(E274G) under its own promoter expedites aggregation of Aβ and α-syn, and drastically reduces their toxicity. A deficiency in AMPylation decreases the cellular tolerance for aggregation-prone polyQ proteins and alters their aggregation behavior. Over-expression of Fic-1(E274G) interferes with cell survival and larval development, underscoring the need for tight control of AMPylase activity *in vivo*. We thus define a link between HSP70 AMPylation and the dynamics of protein aggregation in neurodegenerative disease models. Our results are consistent with a cyto-protective, rather than a cytotoxic role for such protein aggregates.

## Introduction

Neurodegenerative diseases (ND) such as Alzheimer’s, Huntington’s and Parkinson’s are diseases of aging that share a common hallmark: protein aggregates. In healthy cells, chaperones and other heat shock proteins (HSPs) work to maintain protein homeostasis (proteostasis) by reducing the burden of unfolded proteins and overseeing the formation, triage and degradation of protein aggregates. In particular, the HSP70 family of chaperones populates some of the most critical nodes in the proteostasis network, and as such, their expression level and activity is highly regulated. [1–4]. Because of their ability to modulate protein aggregation, several HSPs were suggested to play key roles in the development and progression of NDs. These often involve clogging of the target cells and tissues with accumulations of peptides or proteins such as α-synuclein (α-syn) in Parkinson’s disease, mutant huntingtin (mHtt), which contains extended poly-glutamine (polyQ) repeats, in Huntington’s disease (HD) and amyloid-β-peptide (Aβ) in Alzheimer’s disease (AD) [5–9]. The unifying theme that connects these NDs is their strong association with a failure of cellular maintenance mechanisms to control pathological protein aggregation. Inside cells, such disease-associated, aggregation-prone proteins start out as monomers, only then to participate in a continuous conversion into soluble toxic oligomers. The sequestration of these toxic oligomers into large, insoluble aggregates and inclusion bodies is part of a cellular coping mechanism that gradually depletes these toxic intermediate species of α-syn, Aβ and mHtt [10–16]. Molecular chaperones (e.g. HSP40, HSP70) and chaperonins (e.g. TRiC) orchestrate the continuum of mHtt, Aβ and α-syn oligomerization and alleviate associated cytotoxicity, by preventing monomers from oligomerizing, by facilitating degradation of oligomers via the ubiquitin-proteasome and the autophagy-lysosomal pathways or through enhancement of oligomer deposition into large, insoluble aggregates [17–19]. Because of their prominent role in the modulation of protein aggregation, the master transcriptional regulator of the HSR, HSF-1, and its downstream targets, HSP70 and HSP90 are considered prime targets for intervention in ND [20–22]. However, while up-regulation of HSPs may be beneficial in the context of ND, excessive activity of HSPs favors fast, uncontrolled cell division cycles and is suspected to be a contributing factor to cancer [23–26].

Protein AMPylation is a newly discovered mechanism that regulates HSP70 activity in the ER and the cytoplasm [27–32]. The addition of AMP to the side-chain of a threonine or serine residue is catalyzed by conserved enzymes (AMPylases) that are present in a single copy in most metazoans, including *Caenerhobdatis elegans* (Fic-1), *Mus musculus* (mFICD) and humans (HYPE), but is absent from yeast. HYPE preferentially AMPylates the ER-resident HSP70 family chaperone Grp78/BiP in its substrate-free, ATP-bound conformation [30]. BiP AMPylation disfavors co-chaperone-dependent ATP hydrolysis, believed to be a prerequisite for client binding. AMPylation ‘locks’ BiP in a primed state [33, 34], ready to engage in client refolding immediately after BiP’s de-AMPylation [35]. The *C. elegans* HYPE ortholog Fic-1 modifies a number of ER and cytoplasmic targets, including the HSP70 family members HSP-1 (cytosolic) as well as HSP-3 and HSP-4 (the two *C. elegans* Grp78/BiP orthologs) [29]. AMPylase-deficient fic-1(n5823) animals show enhanced susceptibility to *Pseudomonas aeruginosa* infections, while worms that express a mutant AMPylase with enhanced activity under the control of the endogenous fic-1 promotor *(nIs733* [P*fic-1*::Fic-1(E274G)) display increased pathogen tolerance, suggesting a link between HSP AMPylation, innate immunity and stress tolerance *in vivo* [29]. Expression of active Fic-1(E274G) in yeast – a eukaryote that lacks endogenous AMPylation enzymes – induces massive protein aggregation, which was alleviated by the over-expression of Ssa2, a cytosolic HSP70 protein [27]. These data indiciate a novel mode of HSP70 inactivation by AMPylation and point towards a broad role for protein AMPylation in the regulation of proteostasis.

Here we explored the implication of HSP70 AMPylation in protein aggregation behavior and toxicity in *C. elegans* models of neuro-degenerative diseases. We find that changes in AMPylation levels cause altered aggregation dynamics *in vitro* and *in vivo*, with beneficial or detrimental outcomes, depending on the ND model examined. Expression of active Fic-1(E274G) significantly increases survival of Aβ-expressing worms despite enhanced aggregate formation. RNAi-mediated ablation of HSP-1, HSP-3 and HSP-4 phenocopies these results. Increased AMPylation also alleviates α-syn toxicity. Conversely, we find that expression of aggregation-prone polyQ-proteins in an AMPylase-deficient *fic-1(n5823)* background worsens associated symptoms. Our work indicates a role for HSP70 AMPylation in the control of protein aggregation *in vivo* and highlights the potential of HSP70 as a target for modulation of pathological protein aggregation.

## Results

### Changes in cellular AMPylation levels alter Aβ aggregation and toxicity

AMPylation of HSP70 family proteins prevents them from participating in protein quality control [30]. As such, we hypothesized that intracellular AMPylation levels could affect protein aggregation and associated toxicity of the Alzheimer’s disease-associated (Aβ) peptide, by reducing or enhancing the available pool of active HSP70s. To test this, we mated *Caenorhabditis elegans* strains with constitutive muscular expression of Aβ (CL2006) with previously established AMPylase-deficient *fic-1(n5823)* worms or integrated transgenic nematode strains that express constitutive-active Fic-1(E274G) under the control of the endogenous fic-1 promotor and analyzed homozygous F2 offspring [29, 36]. When transferred from permissive (15 °C) to inducing conditions (20 °C), Aβ aggregation in *C. elegans* body wall muscle cells causes paralysis and significantly shortens life-span [36]. The presence of Fic-1(E274G) was sufficient to alleviate Aβ toxicity under inducing conditions, while AMPylation deficiency did not alter survival or impaired mobility of Aβ-expressing animals (Fig. 1A, Fig. S1A). Analysis of total Aβ content confirmed that Aβ levels remained unaltered in response to changes in AMPylation (Fig. S1B). Because we previously identified HSP70 family members HSP-1, HSP-3 and HSP-4 as prime targets for Fic-1 in *C. elegans*, we asked whether RNAi-mediated reduction of HSP-1, HSP-3, HSP-4 or combinations thereof would suppress Aβ toxicity. While individual knockdowns of HSP-1, HSP-3 and HSP-4 enhanced survival of Aβ-expressing worms only slightly, combined knock-down of HSP-1, HSP-3 and HSP-4 in pairs significantly improved survival, with a triple knock-down of HSP-1, HSP-3 and HSP-4 resulting in ~80 % survival after 4 days at 22 °C, compared to approximately ~20 % survival in controls (Fig. 1B). Similarly, the introduction of an hsp-3 null allele into Aβ-expressing strains ameliorated its fitness, consistent with a beneficial effect of reduced HSP-3 function (Fig. S1C). Ablation of hsf-1 or daf-16, two transcription factors implicated in the control of HSP transcription [37, 38], did not reduce the beneficial effect of Fic-1(E274G) expression in Aβ worms. This result is consistent with the notion that Fic-1(E274G) acts on pre-existing chaperone pools, rather than change the overall abundance of HSF-1- or Daf-16-dependent HSPs (Fig. S1D).

**Figure 1:**
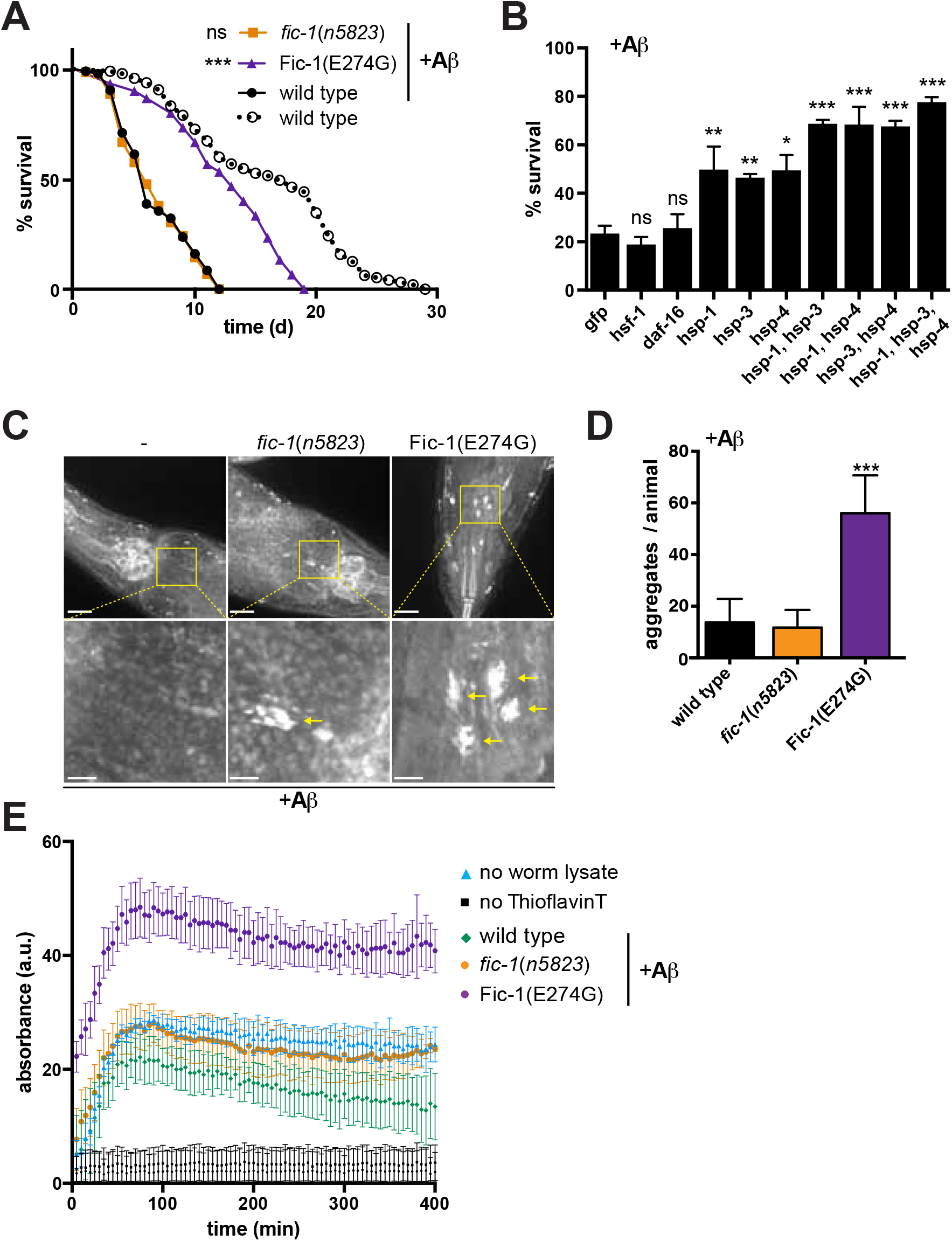
Changes in cellular AMPylation levels alter Aβ aggregation and toxicity. (A) lifespan analysis of wild type, *fic-1(n5823)* and Fic-1(E274G) animals expressing human Aβ peptide from a minigene. (B) Survival assays of wild type animals expressing human Aβ peptide upon RNAi-mediated ablation of indicated genes (x-axis). Thioflavin S stainings (C) and quantification of protein aggregates (D) of wild type (n=31), *fic-1(n5823)* (n=34) and Fic-1(E274G) (n=23) animals expressing human Aβ peptide. Lower panel shows higher magnification images of the boxed areas. Scale bar equals 10 μm (upper panel) and 2 μm (lower panel). For (A) to (D): Error bars represent SD. Statistical significance (P-values) were calculated using the Gehan-Breslow-Wilcoxon test (A) or Mann-Whitney test (B, D) as compared to N2 wild type control (A, D) or anti-GFP RNAi control (B). **\*p < 0.05, **p < 0.01, ***p < 0.001, not significant (ns) p > 0.05.**

To test whether enhanced survival of Fic-1(E274G) worms was accompanied by changes in Aβ aggregation, we stained worms grown under inducing conditions with Thioflavin S to visualize amyloid plaque formation. Fic-1(E274G) strains contained significantly more aggregates than wild-type and *fic-1(n5823)* worms. (Fig. 1C – D). *In vitro* aggregation of Aβ in the presence of *post debris* worm supernatants of Fic-1(E274G) strains was enhanced in comparison with *fic-1(n5823)* or N2 wild-type controls (Fig. 1E). AMPylation of HSP-1, HSP-3 and HSP-4 may thus limit their ability to prevent Aβ aggregation pushing the balance away from toxic intermediate oligomers and toward the formation of large, cyto-protective over intermediate, toxic Aβ aggregates.

### AMPylation deficiency enhances survival under heat stress conditions

When over-expressed in *Saccharomyces cerevisiae*, an organism that lacks endogenous AMPylase activity, the constitutively active *Caenorhabditis elegans* AMPylase Fic-1(E274G) transfers AMP to the yeast cytosolic HSP70 family protein Ssa2 [27]. AMPylation-mediated inhibition of Ssa2 induces the heat shock response (HSR), enhances protein aggregation and eventually causes cell death [27]. To determine whether Fic-1 contributes to the HSR in its endogenous environment, we first introduced a transgenic heat-shock reporter (P*hsp-16.2::gfp*) into *fic-1(n5823)* and Fic-1(E274G) worms. Neither in the absence nor following a heat shock (30 minutes at 35 °C) did we observe genotype-dependent changes in reporter activation (Fig. S2A). We next performed RNAseq experiments to compare wild-type, *fic-1(n5823)* and Fic-1(E274G) strains exposed to heat stress or left untreated. In the absence of extrinsic stress, transcription profiles of the tested strains showed no major transcriptional alterations in response to hypo– *(fic-1 n5823)* or hyper (Fic-1(E274G))-AMPylation (Fig. 2A, S2B-C). Worms exposed to acute heat stress, as expected, showed a substantial upregulation of hsf-1-regulated *hsp* genes (hsp-16.2, hsp16.41, hsp-70, etc.), but this was not affected by the individual genotypes of the tested strains (Fig. 2B, S2D-E). However, a small set of genes – in particular those coding for collagens – showed altered transcript levels that correlated with enhanced or decreased protein AMPylation. qPCR experiments confirmed the absence of AMPylation-related changes in HSR-associated gene expression (Fig. 2C) and corroborated a link between protein AMPylation and collagen gene transcription in Fic-1(E274G) and *fic-1(n5823)* strains (Fig. S2F).

**Figure 2:**
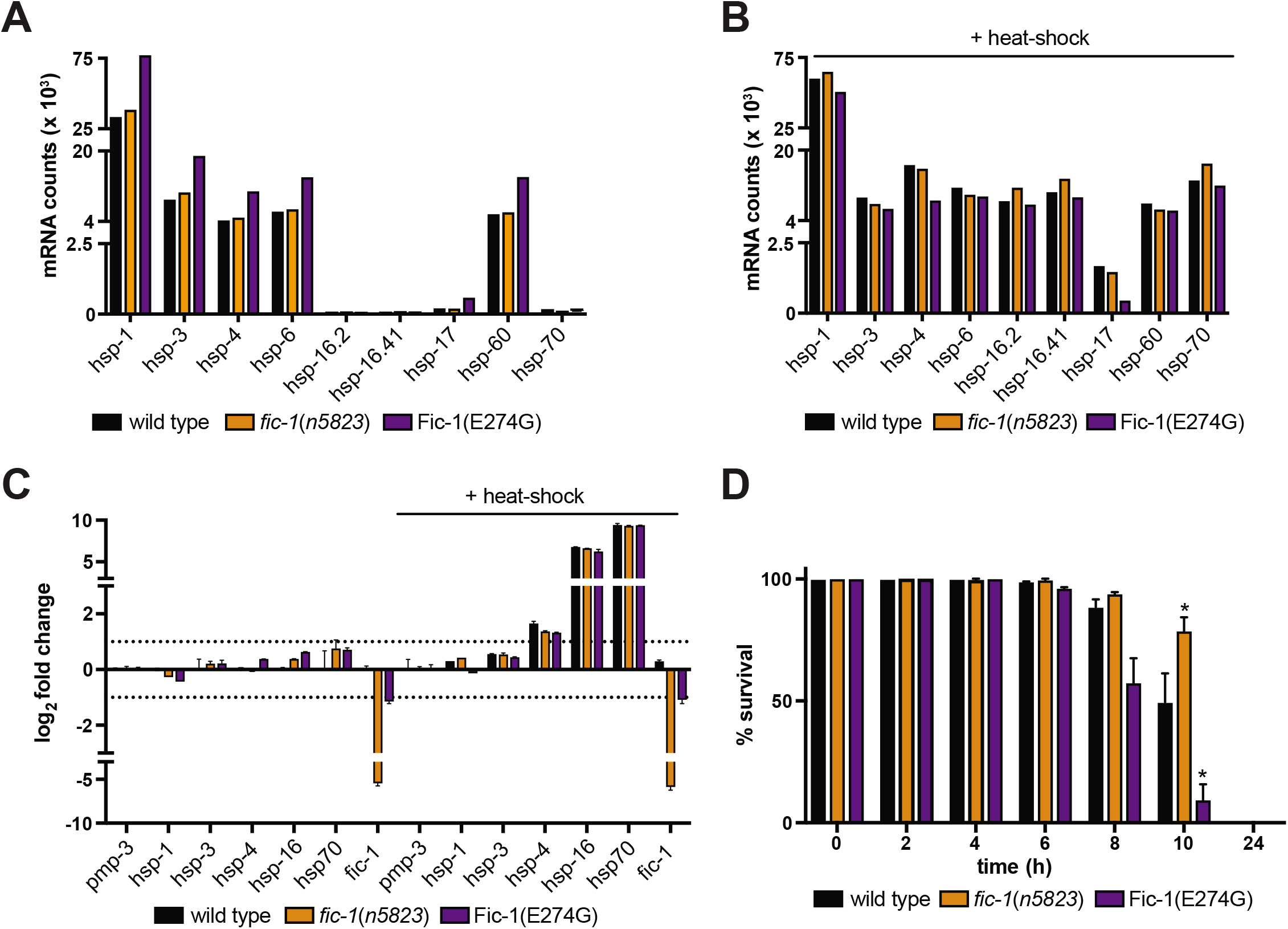
AMPylation deficiency enhances survival under heat stress conditions. (A) heat stress tolerance of wild type, *fic-1(n5823)* and Fic-1(E274G) animals cultured at 33 °C for indicated time points. (B) – (C): normalized mRNA counts of indicated genes in wild type, *fic-1(n5823)* and Fic-1(E274G) animals in the absence (B) or following a 30 minutes heat shock at 35 °C. (D) qPCR validation of transcriptional changes in untreated or heat stressed wild type, *fic-1(n5823)* and Fic-1(E274G) animals. For (A) to (C): Error bars represent SD Statistical significance (P-values) calculated using the Mann-Whitney test compared to N2 wild type control.

To check whether protein AMPylation might play a physiological role in heat stress tolerance beyond induction of the HSR, we tested survival of *fic-1(n5823)* and Fic-1(E274G) worms [29] incubated at 33 °C. Under chronic heat stress (33 °C, 24 hours), *fic-1(n5823)* strains showed enhanced thermo-resistance compared to wild-type controls, while Fic-1(E274G) worms were significantly more sensitive (Fig. 2D). Consistent with this finding, *fic-1(n5823)* embryos survived extended exposure to 33 °C substantially better than wild type or Fic-1(E274G) controls (Fig S2G). In contrast, short-term exposure to heat stress (30 minutes at 35 °C) affected neither worm development nor adult survival after heat-shock in any of the genetic backgrounds tested (Fig. S2H – I).

Together, these results suggest that under prolonged (heat) stress exposure, when the function of HSPs is under constant high demand, HSP AMPylation is critical for survival by imposing an additional layer of HSP regulation that is independent of the transcriptional heat shock response.

### Protein AMPylation affects aggregation behavior of polyQ-repeat containing proteins in *C. elegans*

Transgenic expression of Fic-1(E274G) extended survival of Aβ-positive worms despite enhancing amyloid-β formation. This suggested that changes in intracellular AMPylation levels directly influence the dynamics of protein aggregation *in vivo*. To test this, we asked how aggregation of polyQ-YFP fusion proteins was affected when expressed in *fic-1(n5823)* or Fic-1(E274G) worms. Aggregation of polyQ-repeat variants of YFP in *C. elegans* is age- and polyQ-length-dependent: glutamine repeats of 40 or more residues will promote the formation of discrete, fluorescent foci from early larval stages onwards, while the formation of visible polyQ24-YFP aggregates within 10 days of adulthood is insignificant [39]. Transgenic day 1 adult worms with muscular expression of polyQ40-YFP in a *fic-1(n5823)* background showed a marked increase in the total number of discrete aggregates (45 +/-0.9) when compared to controls (26 +/-0.5) (Fig. 3A) Total levels of polyQ40-YFP as well as transcription of hsp-1, hsp-3, hsp-4 or the hsf-1-regulated HSPs F44H5.4/5 were unchanged (Fig. S3A-B). This distinction, albeit less pronounced with increasing age, persisted on day 5 and day 9 (Fig. 3A, Fig. S3C). The number of aggregates in 9-day-old Fic-1(E274G) worms that express polyQ24-YFP increased significantly (2.9 +/-0.4) compared to controls (0.5 +/-0.1) (Fig. 3B). We also noted a tendency of altered polyQ24-YFP aggregation in AMPylase-deficient *fic-1(n5823)* worms (1 +/-0.2) (Fig. 3B). Consistent with these findings, cellulose acetate filter trap assays of worm lysates corroborated enhanced aggregation of polyQ40-YFP in *fic-1(n5823)* strains, as well as altered polyQ24-YFP aggregation in Fic-1(E274G) and *fic-1(n5823)* strains (Fig. S3D). These findings emphasize the regulatory impact of HSP AMPylation on orchestrating protein aggregation.

**Figure 3:**
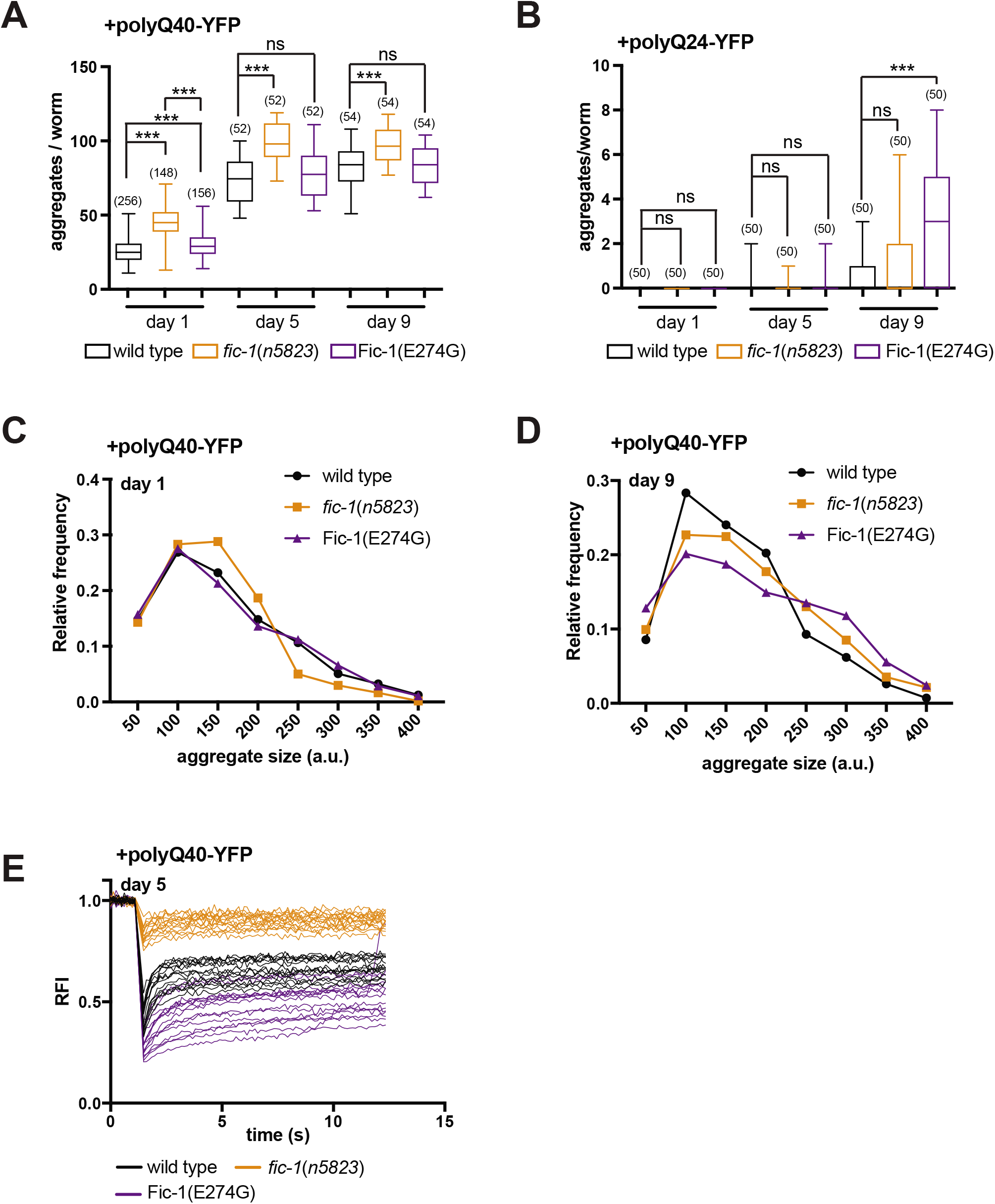
AMPylation controls aggregation of polyQ40 repeat containing proteins in *C. Elegans*. (A-B). Time-dependent aggregation of polyQ40-YFP (A) and polyQ24-YFP (B) in wild type, *fic-1(n5823)* and Fic-1(E274G) animals. The number of tested animals (n) is given in brackets (). Error bars represent SD. Statistical significance (P-value) was calculated using Mann-Whitney test. **\*p < 0.05, not significant (ns) p > 0.05**. (C-D) distribution profile of polyQ40-YFP aggregate sizes on day 1 (C) and day 9 (D) of adulthood. Bin size is 50 a.u. (E) polyQ40-YFP aggregate mobility on day 5 of adulthood as measured by fluorescence after photobleaching (FRAP). Y-axis depicts relative fluorescence intensity (RFI).

Since the presence of Fic-1(E274G) enhanced protein aggregation in Aβ-expressing worms – whereas *fic-1(n5823)* animals showed enhanced polyQ40-YFP aggregate counts – we hypothesized that AMPylation-mediated inhibition of HSPs might change size and mobility of individual polyQ40-YFP aggregates, rather than promote the formation of more foci. We evaluated microscopic images that showed polyQ40-YFP aggregates formed in wild type, *fic-1(n5823)* or Fic-1(E274G) animals and found that on day 1, where the discrepancy between the number of aggregates is maximal, *fic-1(n5823)* worms contained an increased proportion of small aggregates, and a decreased proportion of larger aggregates compared to wild type (Fig. 3C). The presence of Fic-1(E274G) did not alter the overall size distribution of polyQ40-YFP aggregates (Fig. 3C). As worms aged, Fic-1(E274G) expression supported the assembly of larger polyQ40-YFP foci, while the size profile of polyQ40-YFP aggregates in *fic-1(n5823)* strains became more wild type-like (Fig. 3D, S3E). AMPylase deficiency enhanced the mobility of polyQ40-YFP foci, while the presence of Fic-1(E274G) significantly decreased it, as measured by fluorescence recovery after photo bleaching (FRAP) microscopy (Fig. 3E, S3F-G). Changes in intracellular AMPylation levels therefore suffice to modify the abundance-size-mobility relationship of polyQ-repeat protein aggregates.

### AMPylation-mediated changes in polyQ-YFP aggregation alters associated toxicity

Since progressive polyQ-YFP aggregation reduces worm motility and shortens their lifespan [39], we next tested if the severity of the polyQ-YFP-dependent impairments would diverge in a *fic-1(n5823)* or Fic-1(E274G) background. Based on our previous results, we predicted that Fic-1(E274G) strains would exhibit increased fitness relative to wild type or *fic-1* (*n5823*) strains. Indeed, enhanced AMPylation levels significantly extended the lifespan of Fic-1(E274G);polyQ40-YFP animals (Fig. 4A), though we did not observe changes in worm motility over the course of 9 days in relation to AMPylation status or expression of polyQ-protein (Fig. 4B, Fig. S4A) However, polyQ24-YFP-expressing worms seemed particularly sensitive to changes in protein AMPylation, as both Fic-1(E274G) as well as *fic-1(n5823)* significantly shortened lifespan of the animals tested (Fig. S4B).

**Figure 4:**
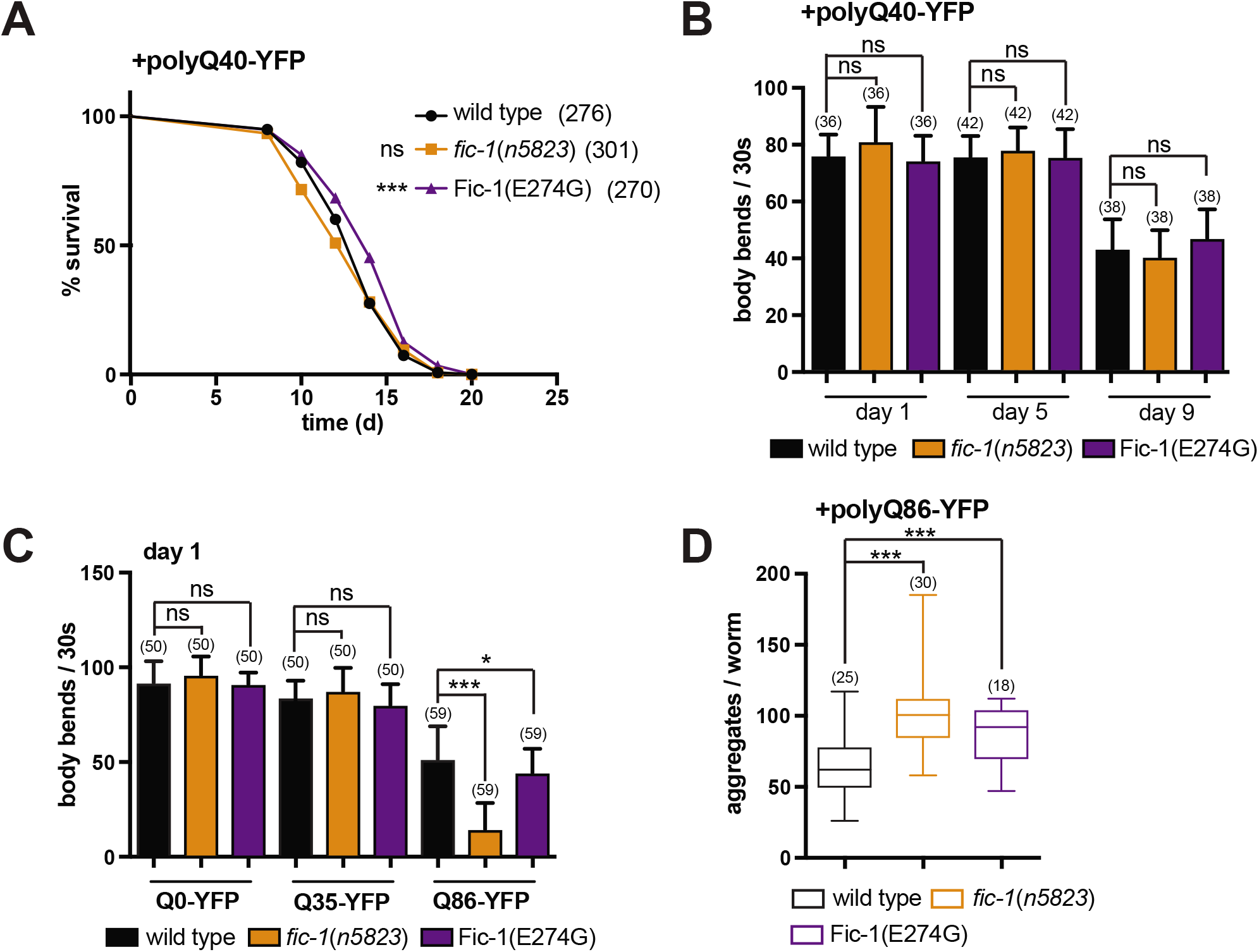
AMPylation affects polyQ-YFP toxicity in *C. elegans*. (A) Motility assay of polyQ40-YFP-expressing wild type, *fic-1(n5823)* and Fic-1(E274G) worms at the indicated time points. (B) lifespan analysis of polyQ40-YFP-expressing wild type, *fic-1(n5823)* and Fic-1(E274G) animals. (C) Motility of wild type, *fic-1(n5823)* and Fic-1(E274G) worms pan-neurally expressing indicated polyQ-YFP proteins at day 1 of adulthood. (D) Number of polyQ86-YFP aggregates in the indicated genomic backgrounds. For (A) to (D): Number of tested animals per group is shown in brackets. Error bars represent SD. Statistical significance (P-values) were calculated using the Mann-Whitney test (motility) or Gehan-Brislow-Wilcoxon test (lifespan) as compared to N2 wild type control. **\*p < 0.05, **p < 0.01, ***p < 0.001, not significant (ns) p > 0.05.**

To evaluate the impact of protein AMPylation on polyQ-YFP aggregation in neurons, we monitored aggregate formation as well as the mobility of transgenic nematode lines with pan-neural polyQ0-YFP, polyQ35-YFP or polyQ86-YFP expression. Loss of Fic-1 potentiated the motility defects (Fig. 4C) and enhanced aggregate formation in polyQ86-YFP-expresing day 1-adults (Fig. 4D, Fig. S4C), but did not affect polyQ0-YFP or polyQ40-YFP strains. Fic-1(E274G) also significantly altered the number of aggregates and motility in polyQ86-YFP expressing strains (Fig. 4C, Fig. 4D).

Together, our data suggest that the AMPylation-mediated regulation of HSPs impacts worm fitness in the presence of of polyQ-repeat proteins, yet to a lesser extent than observed for Aβ-expressing animals.

### AMPylation alters toxicity of α-synuclein aggregates in *C. elegans*

Intrigued by the impact of AMPylation on Aβ and polyQ-YFP toxicity, we asked whether α-synuclein aggregation might be similarly sensitive to changes in Fic-1 activity. We assessed age-dependent α-syn-GFP aggregation in *fic-1(n5823)* or Fic-1(E274G) backgrounds, and found that Fic-1(E274G) expression significantly expedited aggregate formation, such that the number of α-syn-GFP foci in 8-day old α-syn-GFP;Fic-1(E274G) worms matched total numbers of aggregates observed in α-syn-GFP at day 15 of adulthood (Fig. 5A, S5A). Fic-1 deficiency attenuated the assembly of α-syn-GFP foci as worms aged. This yielded significantly fewer aggregates on day 12 or day 15. Consistent with a beneficial role for larger α-syn-GFP aggregates as a means of reducing the toxicity elicited by intermediate species, Fic-1(E274G) worms remained significantly more motile (Fig. 5B) and outlived *fic-1(n5823)* or wild type animals expressing α-syn-GFP in standard lifespan assays (Fig. 5C). While fic-1 deficiency slightly reduced motility (Fig. 5B), it enhanced survival of α-syn-GFP-expressing worms, but not to the extent seen for Fic-1(E274G) animals (Fig. 5C).

**Figure 5:**
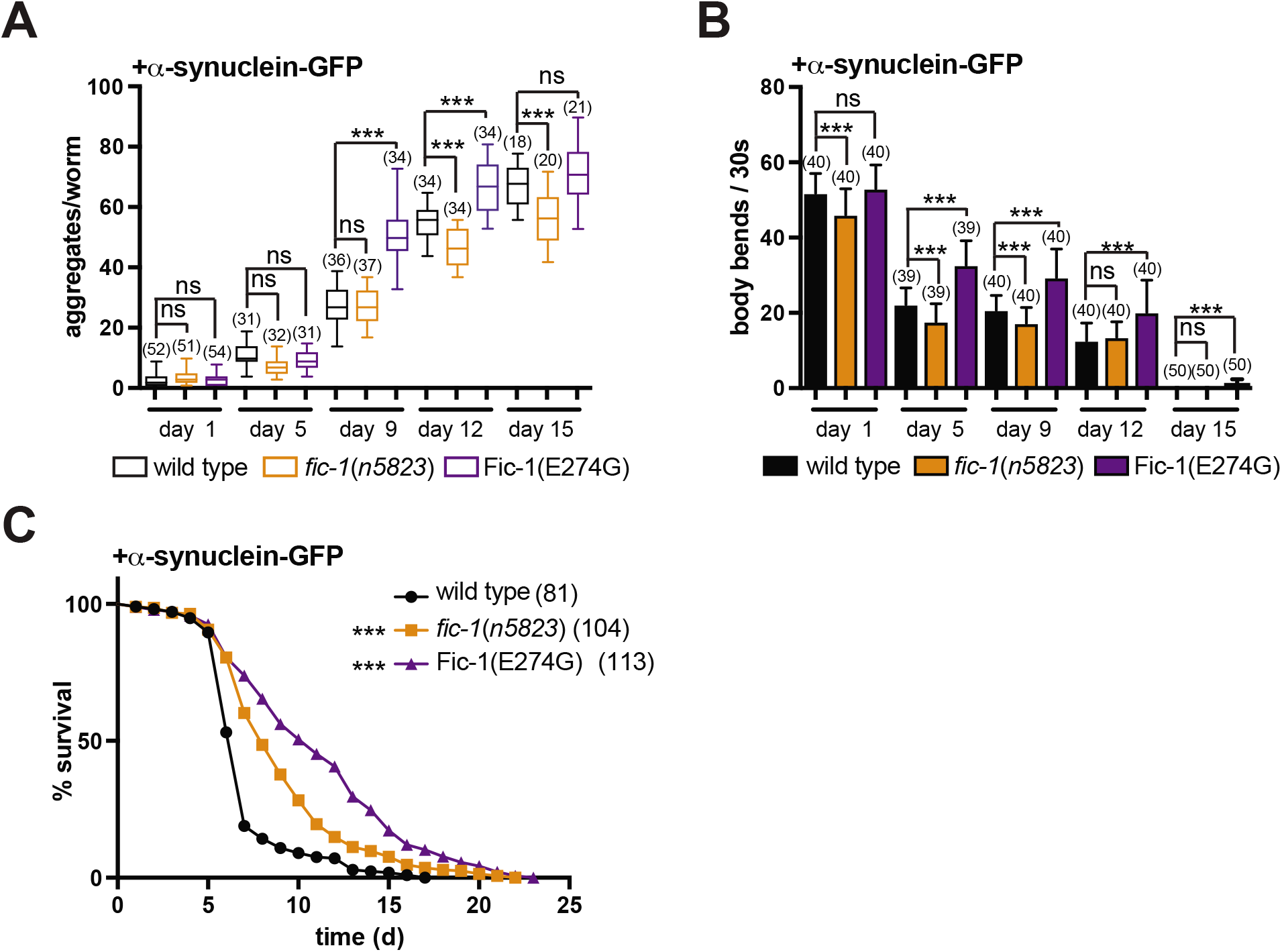
AMPylation alters α-synuclein toxicity in *C. elegans* (A) Time-dependent aggregation of α-synuclein in wild type, *fic-1(n5823)* and Fic-1(E274G) animals. (B) Motility tests of α-synuclein-expressing wild type, *fic-1(n5823)* and Fic-1(E274G) worms at the indicated time points. (C) Survival assay of α-synuclein-expressing wild type, *fic-1(n5823)* and Fic-1(E274G) animals. For (A) through (C): Number of tested animals per group is shown in brackets. Error bars represent SD. Statistical significance (P-values) were calculated using the Mann-Whitney test (motility) or Gehan-Brislow-Wilcoxon test (lifespan) as compared to N2 wild type control. **\*p < 0.05, **p < 0.01, ***p < 0.001, not significant (ns) p > 0.05**.

These data not only support the capacity of fic-1 in the regulation of α-synuclein aggregation, but also indicate additional control of the aggregation-motility-survival relationship beyond protein AMPylation.

### Fic-1 is essential to balance HSP activity during larval development

While endogenous-level Fic-1(E274G) expression does not affect survival of wild type animals [29], it improves worm fitness in the presence of aggregation prone proteins. Because evolution likely favored tight regulation of Fic-1, we suspected that enhanced AMPylation levels could be ambiguous during larval development, where rapid cell division depends on a maximally active proteostasis machinery. We thus asked if AMPylation-mediated inhibition of HSP-1, HSP-3 and HSP-4 could be detrimental during early animal development in worms when proteostasis is challenged by aggregation-prone proteins. Using RNAi, we depleted HSP-1, HSP-3 and HSP-4 in polyQ(40)-YFP and control strains and tested whether larval development was impaired. Whereas development was inhibited upon ablation of HSP-3 or HSP-4 in polyQ40-YFP-expressing wild-type and Fic-1(E274G) L1 larvae, it was unaffected in *fic-1(n5823)* worms (Fig. 6A). Neither HSP-3 nor HSP-4 knock-down affected larval development in the absence of polyQ40-YFP in any of the genetic backgrounds tested, consistent with the previously identified compensatory regulation of the two *C. elegans* BiP orthologs [40]. HSP-1 or combinatorial HSP-3/HSP-4 ablation was lethal at the L1 stage, independent of the presence polyQ(40)-YFP (Fig. 6A). When transferred to RNAi conditions at the L3 stage, knock-down of HSP-1, HSP-3 and HSP-4 did not interfere with aging (Fig. S6A). These results are consistent with an increased demand for chaperone activity to buffer proteostasis in rapidly dividing cells – particularly in the presence of stress caused by protein aggregation – and suggest a regulatory role for Fic-1-mediated HSP AMPylation during early development.

**Figure 6:**
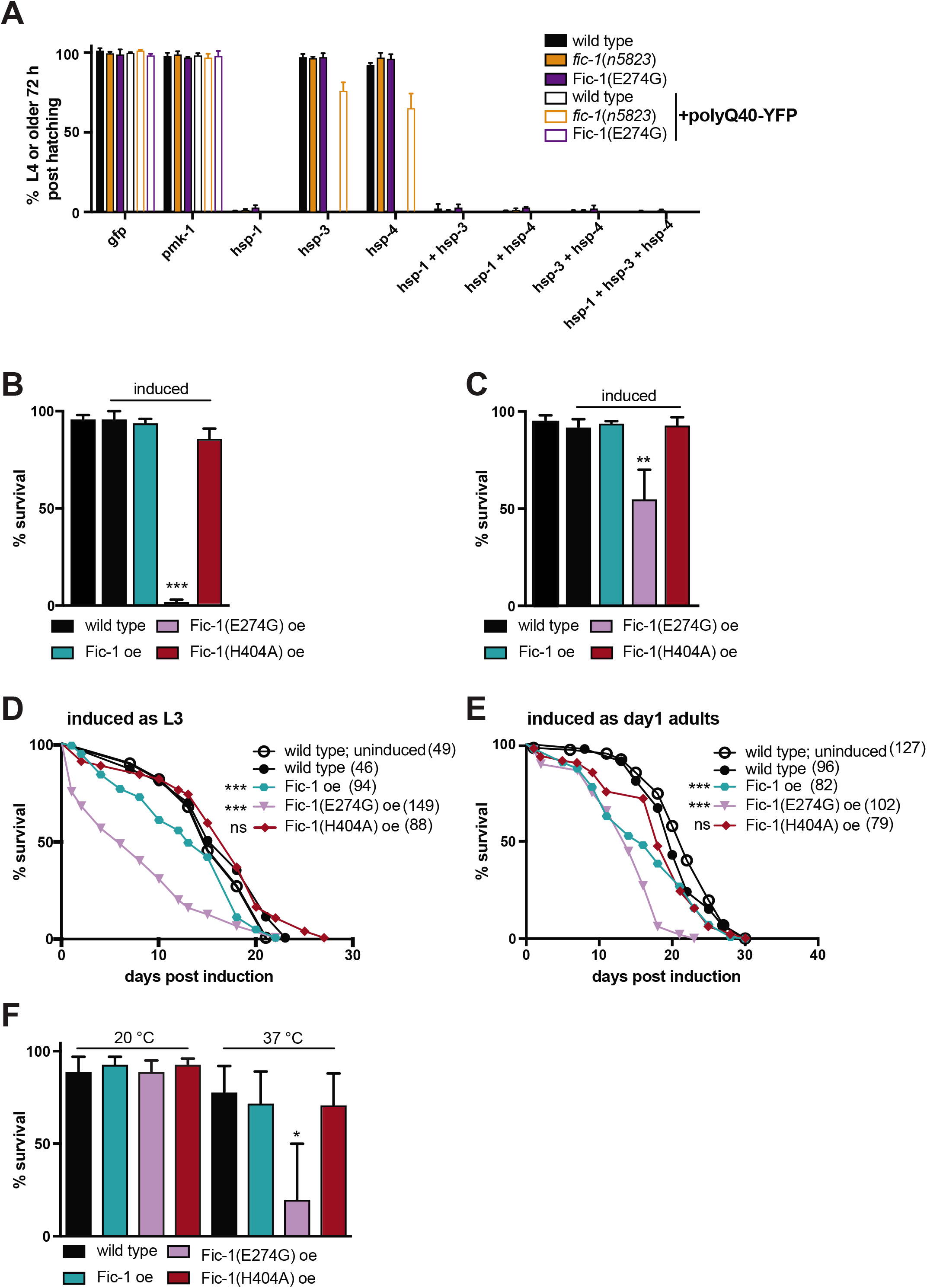
Extensive protein AMPylation interferes with larval development and heat stress tolerance in *C. elegans*. (A) Development assay depicting the proportion of wild type, *fic-1(n5823)* and Fic-1(E274G) embryos that do or do not express polyQ40-YFP and are able to reach adulthood within 72 hours at 20 °C when grown on RNAi plates post-hatching. (B-C) 48 hours survival of L1 (B) or L3 (C) larvae following induction of over-expression (oe) of indicated proteins. (D-E) lifespan assays of worms that over-express the indicated proteins as (D) L3 larvae or (E) day 1 adults. (F) 48 hours survival of day 1 old adults induced to express the indicated proteins. For (A) to (F): Number of tested animals per group is shown in brackets. Error bars represent SD. Statistical significance (P-values) were calculated using the Mann-Whitney test (motility) or Gehan-Brislow-Wilcoxon test (lifespan) as compared to N2 wild type control. **\*p < 0.05, **p < 0.01, ***p < 0.001, not significant (ns) p > 0.05**.

In light of the above, we predicted that HSP hyper-AMPylation might be detrimental in the early larval stages in *C. elegans*. As our attempts to obtain constitutively expressing *Peft-* 3::Fic-1(E274G) strains were unsuccessful, *Peft-3::fic-1* lines were readily obtained. We therefore expressed fic-1, fic-1(E274G) and the catalytically incapacitated mutant enzyme Fic-1(H404A) equipped with a C-terminal HA-tag under the inducible *hsp-16.2p* promotor. If over-expression of Fic-1(E274G) were to lead to systemic HSP70 AMPylation and associated inhibition, then we predicted that the consequences of over-expression would be most pronounced when induced in early larval stages (L1-L2) where rapid cell division is essential. Indeed, we observed an onset-dependent susceptibility to hyper-AMPylation that declined with age: over-expression of Fic-1(E274G) in synchronized L1 larvae was acutely lethal while over-expression of Fic-1 or Fic-1(H404A) did not affect larval fitness or development (Fig. 6B), recapitulating acute toxicity of Fic-1(E274G) previously described in *Saccharomyces cerevisiae* [27]. When induced at the L3 larval stage, Fic-1(E274G) significantly impacted larval development (Fig 6C) and only about 50 % of Fic-1(E274G) worms reached adulthood within 48 hours. The lifespan of this fraction of Fic-1(E274G) animals was significantly reduced when compared to Fic-1 or wild type controls (Fig. 6D). Over-expression of Fic-1(E274G) in day 1 adult worms attenuated survival, yet did not alter motility (Fig. 6E, Fig S6B-C). Notably, over-expression of Fic-1 significantly reduced survival without impairing larval development, possibly because of a higher abundance of endogenous Fic-1 activators in (late) adulthood.

Because AMPylation deficiency conferred enhanced heat tolerance (Fig. 2D), we tested if Fic-1(E274G) over-expression might limit the worm’s ability to sustain prolonged heat stress through inhibition of HSP activity by AMPylation. We induced over-expression of Fic-1, Fic-1(E274G) and Fic-1(H404A), rested the worms for 2 hours at 20 °C to allow protein expression and then exposed them to either 20 °C or 33 °C. Over-expression of Fic-1(E274G) significantly reduced heat tolerance, while it did not affect short-term survival at 20 °C (Fig. 6F).

Our results indicate a particular sensitivity to HSP70 dysregulation in early life and highlight a critical, regulatory role for Fic-1 in balancing cellular HSP activity to accommodate aggregation or heat stress at a stage of rapid cell division.

## Discussion

As cells age, their ability to control the many facets of protein folding declines, producing an environment prone to protein aggregation. Likewise, the cytoprotective HSR and UPR^ER^ are attenuated in older animals [41–43]. HSP70 chaperones are essential in protein (re)folding [44, 45], but whether post-translational HSP70 modifications contribute to the coordination of the cellular chaperoning machinery remained unknown. Our results provide evidence that Fic-1-mediated AMPylation of HSP70 affects the dynamics of protein aggregation by negatively regulating its activity.

Using *C. elegans* models for Aβ, polyQ and α-synuclein aggregation, we found that both reduction and enhancement of intracellular AMPylation alters protein aggregation behavior. Aggregate size and mobility are affected, causing changes in lifespan and motility. We had previously identified HSP-1, HSP-3 and HSP-4 as substrates for the *C. elegans* AMPylase Fic-1 [29]. AMPylation inhibits the activity of HSP-1, HSP-3 and HSP-4, an outcome we recapitulated by their ablation by means of RNAi. We therefore suggest that changes in AMPylation of additional Fic-1 substrates, including the translation elongation factor EEF-1A.2 or core histones [29], may be less important in this context.

AMPylation impairs the activity of HSP-1, HSP-3 and HSP-4, all of which interact with Aβ plaques in *C. elegans* [46], and promotes the formation of larger aggregates. This extends lifespan. Intermediate-sized, soluble protein aggregates are thought to be the major toxic aggregate species, whereas larger, insoluble foci inflict less harm [13-15, 18, 47, 48]. Dampening HSP70 activity through AMPylation, may thus favor assembly of larger foci, as HSP70s are no longer in sufficient supply to counteract protein aggregation (Fig. 7A-B). AMPylation of HSP-3/HSP-4 might impair the turnover of misfolded secretory proteins and enhance the cell’s vulnerability to aggregate formation. Likewise, we propose that AMPylation of HSP-1 – a HSP70 family member that shuttles between the cytoplasm and the nuclear lumen – renders the cytoplasmic and nuclear compartments more vulnerable to protein aggregation.

**Figure 7:**
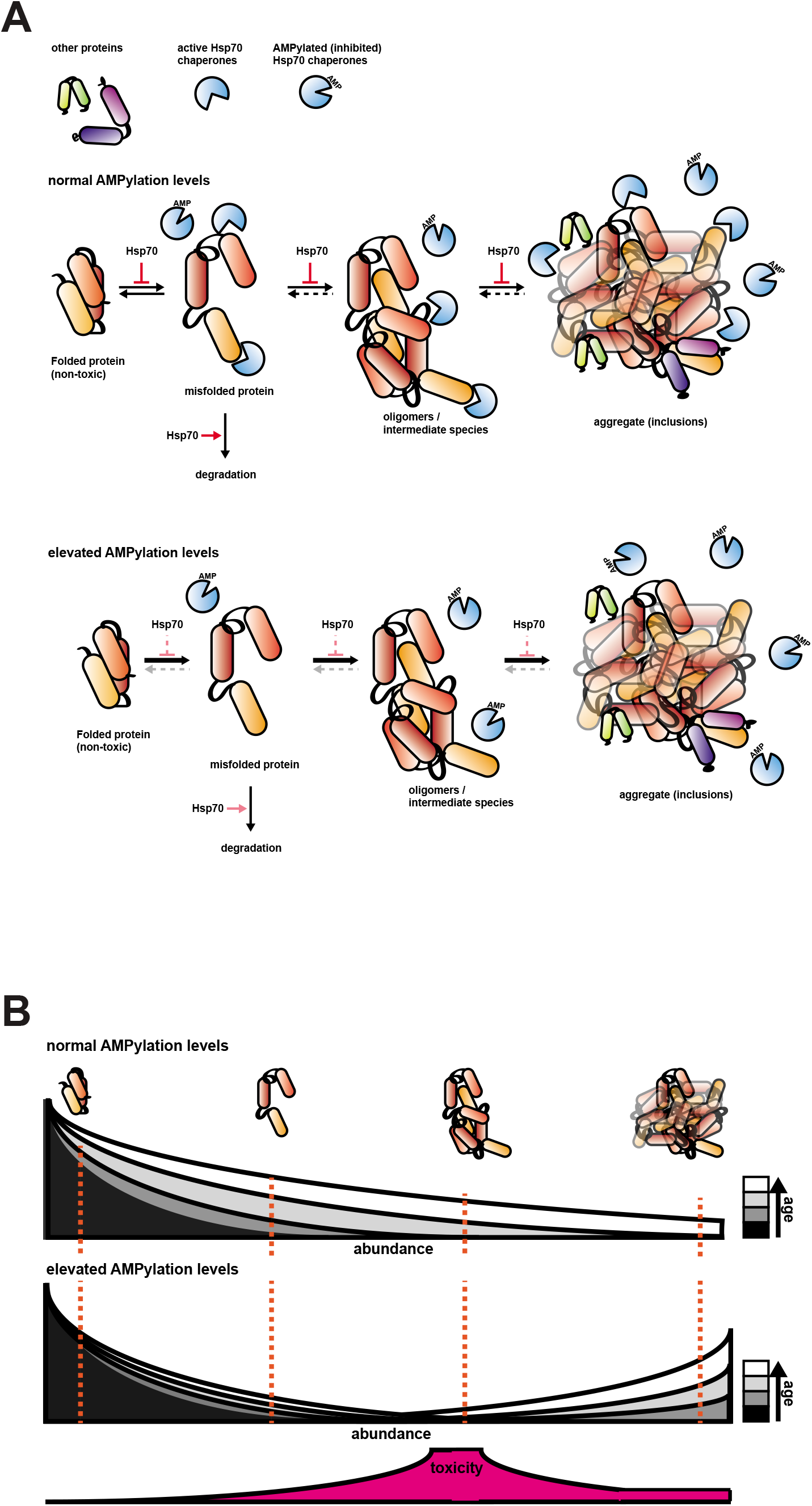
AMPylation levels regulate aggregation behavior. (A) Under normal conditions, HSP70 proteins prevent the unfolding and support the refolding of misfolded proteins, while suppressing the formation of protein oligomers / intermediates as well as aggregates (upper panel). Elevated AMPylation levels limit the available pool of active HSP70 proteins, hence increasing the rate of protein aggregation and favoring the formation of large foci over smaller oligomers (lower panel). (B) Schematic of the aggregation-age – toxicity – AMPylation level relationship.

Connections between HSP70 and Grp78/BiP activity and protein aggregation in mammals are well-established. Several HSP70 activators are currently being tested for their potential to slow the progression of AD, HD and PD [21, 22, 49-52]. Our results suggest that HSP70 inhibitors, rather than activators, might represent an interesting alternative route to explore. What, then, is the possible explanation why both HSP70 activation or inhibition can be advantageous in the context of aggregation-associated toxicity? Under the assumption that soluble, intermediate aggregates represent the major toxic species, both the avoidance of aggregate formation – achieved by HSP over-expression or HSP activators – as well as the expedited assembly of large, cyto-protective aggregates – as stimulated by transient HSP inhibition – would efficiently detoxify affected cells (Fig. 7A-B) [52, 53]. Thus, we propose that pharmacological interventions that target the human AMPylase HYPE may have therapeutic potential.

Enhanced AMPylation levels are beneficial in the context of Aβ and α-synuclein aggregation. For polyQ-repeat proteins, the absence of fic-1, more so than Fic-1(E274G) expression, affects the formation of aggregation foci. The minimal fitness gain seen for polyQ-YFP;Fic-1(E274G) strains suggests that AMPylation levels achieved by endogenous Fic-1 activity might suffice for proteostasis in the presence of this polyQ-YFP species. At this time, however, we do not understand the fundamental mechanisms by which an activating signal for Fic-1 is generated nor how the enzyme perceives it. The impact of HSP AMPylation on protein aggregation and larval development implies a well-balanced regulatory regimen. We further note that Fic-1 expression is elevated in the *C. elegans* germline and in embryos, consistent with an important role of protein AMPylation during larval development [29].

In conclusion, our findings challenge the long-standing paradigm that proteostasis is regulated solely by the abundance of HSP proteins. Rather, a dynamic interplay between expression level and post-translational modifications such as AMPylation coordinate stress tolerance and contribute to age-associated pathologies. Pharmacological modulation of AMPylation may thus provide a new toehold to modulate the proteostasis network for therapeutic benefit in a wide range of age-related diseases.

## Acknowledgements

We thank the members of the Ploegh and Pincus labs for helpful comments and discussion. The Whitehead genome core facility is acknowledged for processing RNAseq and genomic DNA samples. We thank Justin Reemer for technical assistance and the Whitehead FACS facility for support with the reporter assays. George Bell and the Whitehead bioinformatics group are acknowledged for their help with the analysis of RNAseq data. M.C.T. is a recipient of a Young Investigator Award from the Emerald Foundation, Inc. This work was supported by an NIH Early Independence Award (DP5 0D017941-01 to D.P.), and a Cancer Research Fellowship from the Alexander and Margaret Stewart Trust (D.P.).

## Author contributions

HLP supervised the project. MCT and HLP planed and designed the experiments. MCT performed all experiments except for RNAseq data analysis, which was performed by DP. MCT and HLP wrote the manuscript.

## Material and Methods

### *C. elegans* strains and grow conditions

Worms were grown at 20 °C on plates containing nematode growth medium agar seeded with OP50 *E.coli* bacteria [54]. Strains used in this study: N2 Bristol, MT22849 *(n5823* IV *[fic-1* KO]), MT23503 *(nIs733* [*Pfc-1*::Fic-1(E274G), *Pmyo-3::mCherry]), MT22798 (nEx2237 [Pfic-* 1::Fic1(E274G), Pmyo-3::mCherry]), RB1104 *(hsp-3(ok1083))* CL2006 *(dvIs2*[pCL12(P*unc-* 54::human Abeta peptide 1-42 minigene) + pRF4]), MT23526 *(n5823* IV; *dvIs2)*, MT23525 *(nIs733; dvIs2)\**, MT24052 *(hsp-3(ok1083)* X; *dvIs2)*, CL2070 (*dvIs70* [P*hsp-16.2::GFP* + rol-6(su1006)]), MT25130 *(n5823* IV; *dvIs70)*, MT25131 *(nEx2237;dvIs70)*, AM138 *(rmIs130* [P*unc-54*::Q24::YFP]), MT24431 *(n5823* IV; *rmIS130)*, MT24433 *(nIs733; rmIs130)*, AM141 *(rmIs133*[P*unc-54::Q40::YFP])*, MT24058 *(n5823* IV; *rmIS133)*, MT24057 *(nIs733; rmIs133)*, AM1126 *(rmIs383[Pf25b3.3::Q(35)::YFP])*, MT24451 *(n5823* IV, *rmIs383)*, MT24449 *(nIs733, rmIs383)*, AM322 (rmIs135[Pf25b3.3::Q(86)::YFP]), MT25133 *(n5823* IV, *rmIs135)*, MT25132 *(nIs733, rmIs135)*, AM52 (rmIs182[Pf25b3.3::Q(0)::YFP]), MT24446 *(n5823* IV, *rmIs182)*, MT24447 *(nIs733, rmIs182)*, OW40 zgIs15[P*unc-54*::αsyn::YFP], MT24072 *(n5823* IV; *zgIS15* IV), MT24073 *(nEx2237; zgIs15* IV), MT24418 (*nIs776* [P*hsp-16.2::fic-1*, Pmyo-3::mCherry]), MT24419 (*nIs776* [P*hsp*-16.2::Fic-1(E274G), Pmyo-3::mCherry]), MT24420 (*nIs778* [P*hsp-16.2::* Fic-1(H404A), *Pmyo-3*::mCherry])

* strain MT23525 (nIs733; dvIs2) was periodically rebuilt

### Plasmid construction

For inducible protein expression, the *fic-1* gene was cloned into pPD49.78 (Fire lab vector kit) with an additional C-terminal HA-tag introduced as part of the primer to obtain P*hsp-16.2::fic-1::HA*. Plasmids P*hsp-16.2::fic-1 E274G::HA* and P*hsp-16.2::fic-1 E274G::HA* were obtained by site directed mutagenesis of P*hsp-16.2::fic-1::HA* using the Q5 site-directed mutagenesis kit (NEB). P*unc-54::fic-1::mCherry* was assembled in a Gibson reaction containing KpnI & SacI-digested pPD30.38 (Fire lab vector kit) as well as fic-1 and mCherry extended with appropriate homology overhangs. Table S1 depicts all primers used in this study.

### Germline transformation

Inducible *fic-1* overexpression constructs were injected at 50 ng/μl into N2 wild type animals with 10 ng/μl each of *Pmyo-3::mCherry* and *Prab-3::mCherry* as a co-injection markers. P*unc-54::fic-1::mCherry* was injected at 50 ng/μl into N2 wild type with 50 ng/μl of insert-free pcDNA3.1 plasmid.

### Worm synchronization

Asynchronous worm populations were washed off NGM plates using H_2_O and collected in 1.5 ml tubes. Worms were centrifuged at 4200 rpm for 30 seconds and H_2_O was aspired. Worms were lysed in 1 ml of hypochlorite bleaching buffer for exactly 7 minutes with shaking (1200 rpm) on a heat block set to 20 °C and treatment-resistant embryos were recovered by centrifugation at 4200 rpm for 30 seconds. Worms were washed twice with 1 ml of H_2_O and embryos were thereafter transferred to fresh OP50-seeded NGM plates. To avoid unwanted progressive selection of bleach-resistant worms through repetitive hypochlorite treatment, following synchronization, worms were used only for the intended immediate experiment and discarded thereafter. Worm stock plates were explicitly spared from hypochlorite treatments.

### In vivo Aβ aggregation assay

For paralysis and lethality tests, L3 larvae from strain CL2006 or derivatives thereof were transferred to fresh NGM plates containing an *E.coli* OP50 lawn and incubated at 20 °C. Animals were inspected at the indicated intervals, but at least once a day and scored according to their fitness. At each time point, an animal would be assigned to one of three categories: unaffected (no apparent movement impairment), paralyzed (major body axis paralysis with remaining, limited mobility of head and tail tip) or dead (full-body paralysis and irresponsive to at least 10 pokes with a platinum loop and soft tickling using a human hair)

### In vivo polyQ-YFP and α-syn-GFP aggregation assays

Assays were adapted from [39]. In brief, synchronized animals expressing indicated polyQ-YFP or α-syn-GFP species were grown on NGM plates containing an *E.coli* OP50 lawn. Total aggregates were counted at the indicated time points by visual inspection, using a stereomicroscope equipped for epifluorescence. Aggregates were defined as discrete, stand-alone structures visibly separated from the surrounding fluorescence. For bouh polyQ-YFP or α-syn-GFP-expressing strains, the variance in aggregate counts of a single animal scored multiple times was less than 15 %. Relative changes in aggregate numbers (e.g. *fic-1(n5823)* animals contain visibly more aggregates than wild type) were confirmed by at least one double-blinded observer.

For polyQ aggregate size measurements, animals were immobilized on a 1 % agar pad using 0.2 % levamisole and imaged using a 20x objective. At least 20 images per strain depicting at least 20 randomly chosen worm segments were analyzed using Fiji to semi-automatically retrieve aggregate numbers, their intensities and size.

### RNA interference

RNAi feeding was performed as described [55]. NGM plates supplemented with 1 mM IPTG were spotted with HT115 bacteria containing a L4440 plasmid derivative targeting the respective genes of interest; L4440 encoding for either a anti-*bam-1* or anti-*gfp* RNA were used as mock controls. Embryos or larvae were transferred at the indicated time points to RNAi plates and incubated at 20 °C unless state otherwise. For combinatorial RNAi experiments, similar amounts of each individual HT115-RNAi *E.coli* strain were spotted onto IPTG plates.

### Motility assays

Synchronized worm populations were incubated at 20 °C and assessed for motility at the indicated time points, following a protocol described in [56]. Briefly, animals were transferred to a 50 μl droplet of M9 on a glass slide and allowed to acclimatize for 60 seconds. Thereafter, the number of full body bends was assessed for 30 seconds. Each experiment was independently performed three times with at least 10 animals scored per experiment.

### Thioflavin S staining of amyloid plaques and microscopy

Aβ aggregates were stained as describe in [57]. In brief, CL2006 nematodes were fixed in 4 % paraformaldehyde/PBS, pH 7.5, for 12 hours at 4 °C, permeabilized in 5 % fresh β-mercaptoethanol, 1 % Triton X-100, 125 mm Tris, pH 7.5, at 37 °C for 24 hours and stained with 0.125% thioflavin S (Sigma) in 50 % ethanol for 2 min. Animals were destained in 50 % EtOH for 2min, washed 3x with ddH_2_O and transferred in approximately 10 μl volume on a droplet of Fluoromount G on a glass slide for microscopy. All images were collected on a PerkinElmer Ultraview Multispectral Spinning Disk Confocal Microscope equipped with a Zeiss 1.4 NA oil immersion 63x objective lens and a Prior piezo-electric objective focusing device. Images were acquired with a Hamamatsu ORCA ER-cooled CCD camera controlled with Metamorph software. Post-acquisition image manipulations were made using Fiji software [58]. For analysis of amyloid plaques, at least 20 randomly chosen CL2006 nematodes per replica were assessed by eye and the number of aggregates counted.

### Immunoblotting

Nematodes were washed off NGM plates, collected by centrifugation and resuspended in 50 mM Tris-HCl pH 8.0, 150 mM NaCl, 5 mM EDTA, 1 % NP-40, 0.1% SDS. Samples were homogenized using a Qiagen TissueLyser II bead mill (30 Hz, 5 minutes), centrifuged for 1 minute at 12’000 g to pellet worm debris, supplemented with SDS sample buffer and subjected to SDS-PAGE. Protein transfer and antibody probing was performed as described in [29]. Antibodies used in this study are listed in table S3.

### Development and stress assays

Embryos obtained by hypochlorite treatment of asynchronous worm populations were transferred onto assay plates and the number of embryos per plate was counted. Unless stated otherwise, animal development was scored by visual inspection after 96 hours of incubation at 20 °C. Animals that failed to reach the L4 larval stage were considered developmentally defective.

For heat tolerance assays, approximately 100 animals per plate were incubated at the indicated temperatures and scored at regular intervals. Animals irresponsive to repetitive (10x) poking with a platinum loop and unable to recover within 60 min at 20 °C were considered dead. Experiments were terminated after the last animal died.

### Fitness and longevity assays

Day1 adults were placed onto fresh NGM plates and then transferred onto new NGM plates at two-day intervals throughout the experiment. We explicitly avoided the use of FUDR as we observed beneficial effects of FUDR on Aβ tolerance. Aged animals were repeatedly poked (up to 10x) with a platinum loop and scored. Animals were considered dead if repetitive poking (10x) did not result in any visible body movement. Dead animals were immediately removed from assay plates.

### RNAseq andRT-qPCR

Synchronized worm populations kept on 2-5 10 cm2 NGM-OP50 plates were grown to day 1 of adulthood at 20 C and either heat-shocked at 35 C for 2 hours followed by 60 minutes of recovery at 20 C or directly collected. Worms were washed 6x with 1 ml H2O and 3x with RNAse-free H2O and snap-frozen in liquid nitrogen. Worms were resuspended in RLT Plus buffer and lysed using a Qiagen TissueLyser II bead mill (30 Hz, 5 minutes). Total RNA was extracted using RNeasy Mini kit (Qiagen) and submitted to the Whitehead Genome Technology Core where samples were processed as described in [27]. Reads were mapped using Tophat and quantified using DEseq2.

For RT-qPCR, total RNA was reverse transcribed using the SuperScript III first strand synthesis system (Invitrogen) and analyzed on a QuantStudio 6 flex real time PCR system. Table S2 lists all primers used in RT-qPCR tests.

### In vitro aggregation assays

*In vitro* Aβ aggregation assays were performed as described in [59]. In brief, worms were collected, resuspended in 50 mM Tris-HCl pH 8.0, 150 mM NaCl, lysed by sonication and centrifuged to pellet *C. elegans* debris. Cleared worm lysates were supplemented with recombinant Aβ_1-42_ (10 μM final concentration) and Thioflavin T (20 μM final concentration) in a total volume of 100 μl. Samples were transferred into a black 96-well microplate and incubated at 37 °C. Thioflavin T fluorescence was measured at 10 minutes intervals for 24 hours (excitation at 440 nm, emission at 485 nm) using a SpectraMax M3 plate reader (Molecular Devices).

### Fluorescence Recovery after Photobleaching (FRAP)

For FRAP analysis of aggregate mobility, worms were mounted on agar pads and immobilized with 0.2 % levamisole. Animals were analyzed using a Zeiss LSM710 Laser Scanning Confocal microscope equipped with a Zeiss AxioObserver (motorized and inverted microscope stand with DIC optics), a 63x oil objective lens. The hardware was controlled by the Zeiss Zen Balck 2012 acquisition software. FRAP settings were adapted from [60]. In brief, 0.623 μm^2^ were bleached (35 iterations, 100% transmission) and images were collected at 10 ms intervals. Relative fluorescence intensity (RFI) was determined as described in [61].

### Experiment statistics

Statistical analyses were performed in Graphpad Prism (GraphPad Software). The individual statistical tests utilized in each experiment as well as number of worms per group are indicated in the respective figures and figure legends. Unless indicated otherwise, Mann-Whitney tests (non-parametric t-test that compares ranks of unpaired samples) were performed with N2 wild type levels serving as baseline. Lifespan assays were probed employing the Gehan-Brislow-Wilcoxon test.

---

**Figure S1:**
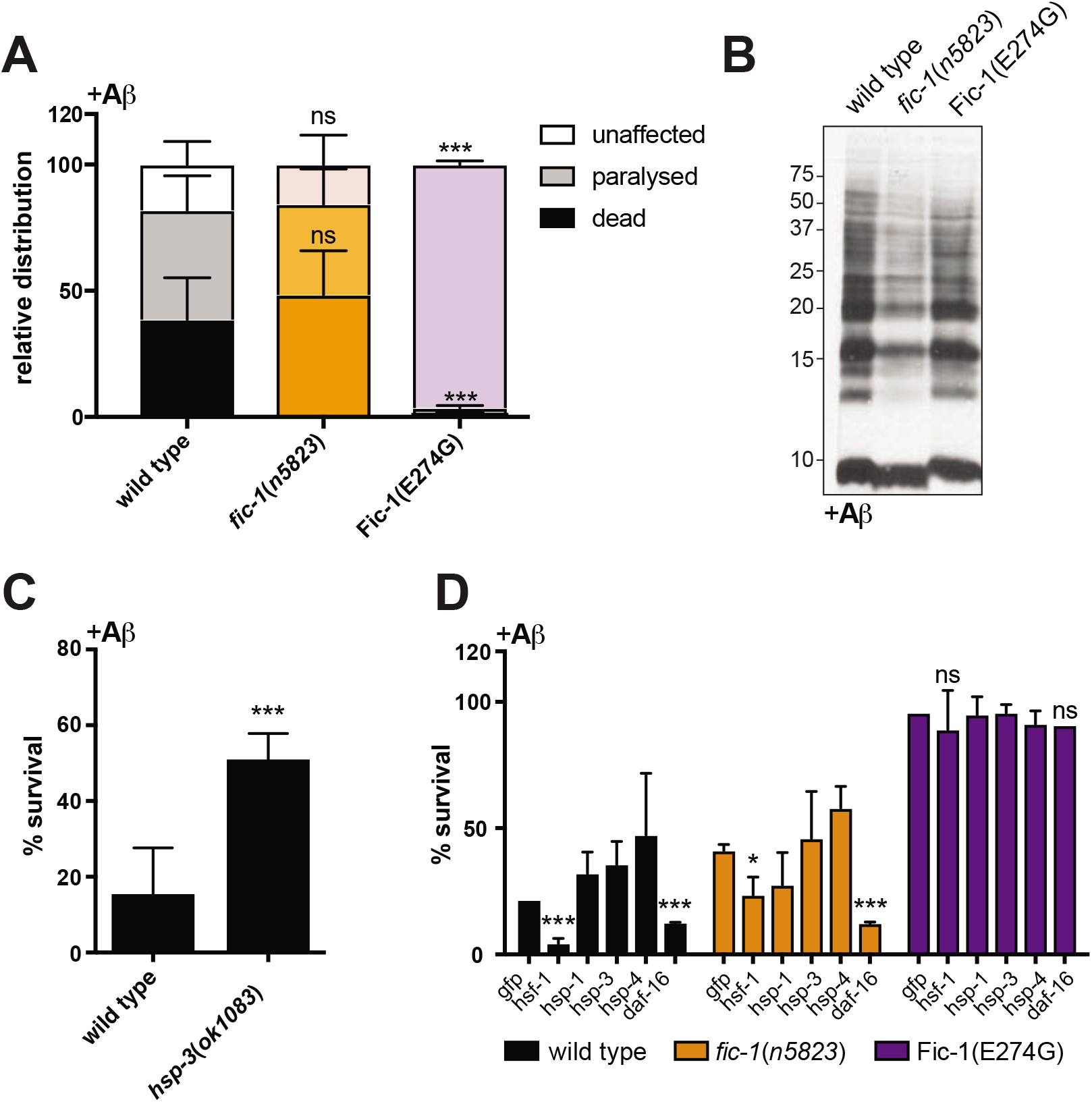
Changes in cellular AMPylation levels alter Aβ aggregation and toxicity. (A) Fitness analysis of wild type, *fic-1(n5823)* and Fic-1(E274G) animals expressing human Aβ peptide from a minigene after 96 hours of incubation at 20 °C. Paralyzed refers to animals shwoing major body axis paralysis with remaining minor mobility of tail tips and heads. (B) Immunoblot of indicated *C. elegans* total lysates. Total protein concentrations were determined by a Micro BCA assay and equal amounts of protein were loaded in each lane. Aβ was detected using anti Aβ Ab clone 6E10. (C) Survival of N2 and *hsp-3(ok1083)* animals expressing human Aβ peptide. (D) Survival assays of wild type, *fic-1(n5823)* and Fic-1(E274G) animals expressing human Aβ peptide upon RNAi-mediated ablation of indicated genes (x-axis). For (A) to (D): Error bars represent SD. Statistical significance (P-values) were calculated using the Mann-Whitney test as compared to N2 wild type controls. **\*p < 0.05, **p < 0.01, ***p < 0.001, not significant (ns) p > 0.05**.

**Figure S2:**
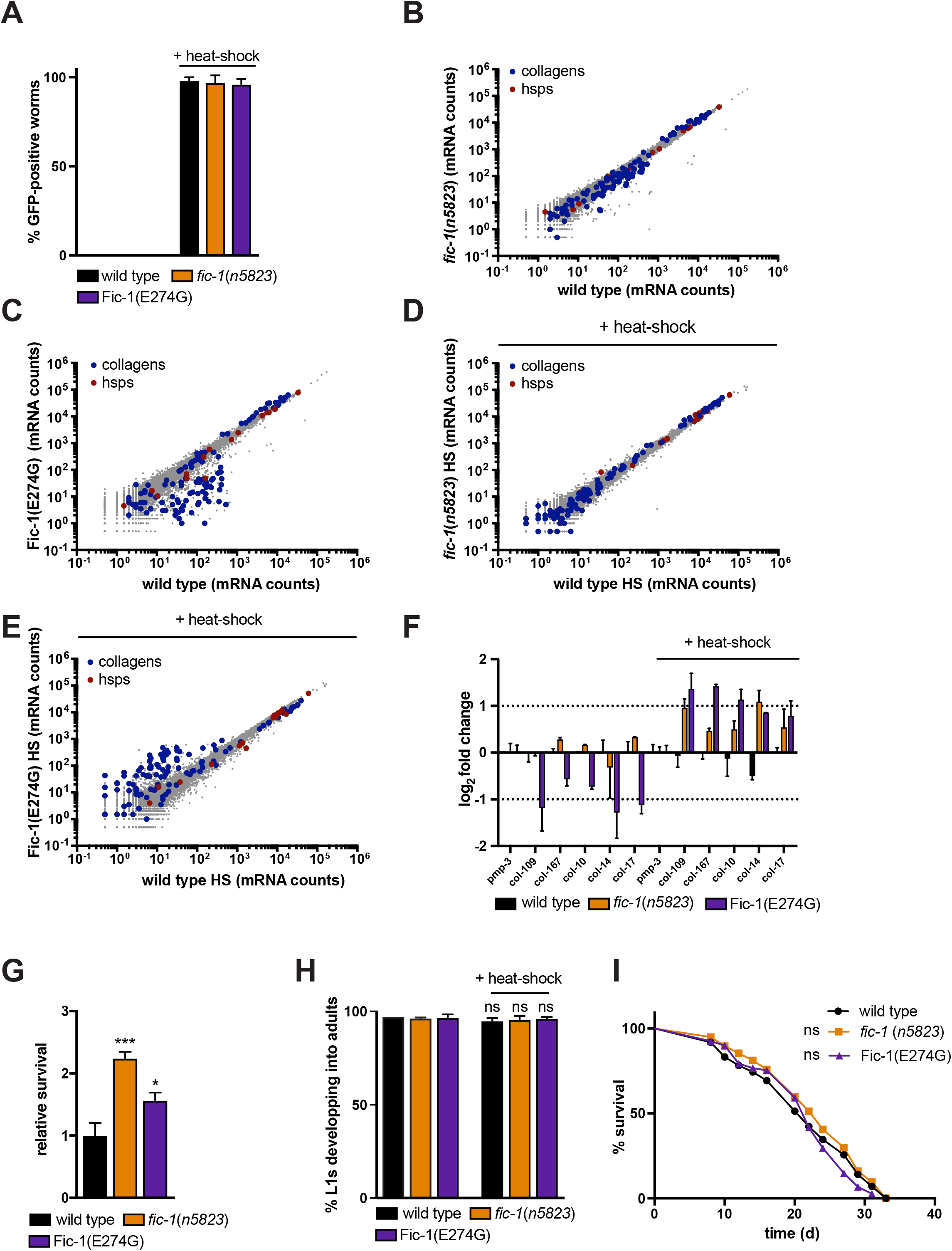
AMPylation deficiency enhances survival under heat stress conditions. (A) Larvae to adult development of indicated strains following heat-shock for 60 minutes at 35 °C and 48 hours of incubation at 20 °C. (B) survival curve of wild type, *fic-1(n5823)* and Fic-1(E274G) animals heat-shocked for 30 minutes at 35 °C and thereafter incubated at 20 °C. Error bars represent SD. Statistical significance (P-values) were calculated using the Gehan-Breslow-Wilcoxon test as compared to N2 wild type control. **\*p < 0.05, **p < 0.01, ***p < 0.001, not significant (ns) p > 0.05**. (C) – (F): Global changes in transcript levels comparing *fic-1(n5823)* or Fic-1(E274G) to wild type animals. Worms were cultured at 20 °C and collected prior ((C), (D)) or after ((E), (F)) exposure to acute heat stress (30 minutes at 35 °C) Heat shock protein genes (hsps) are highlighted in red, collagen genes (collagens) are colored in blue.

**Figure S3:**
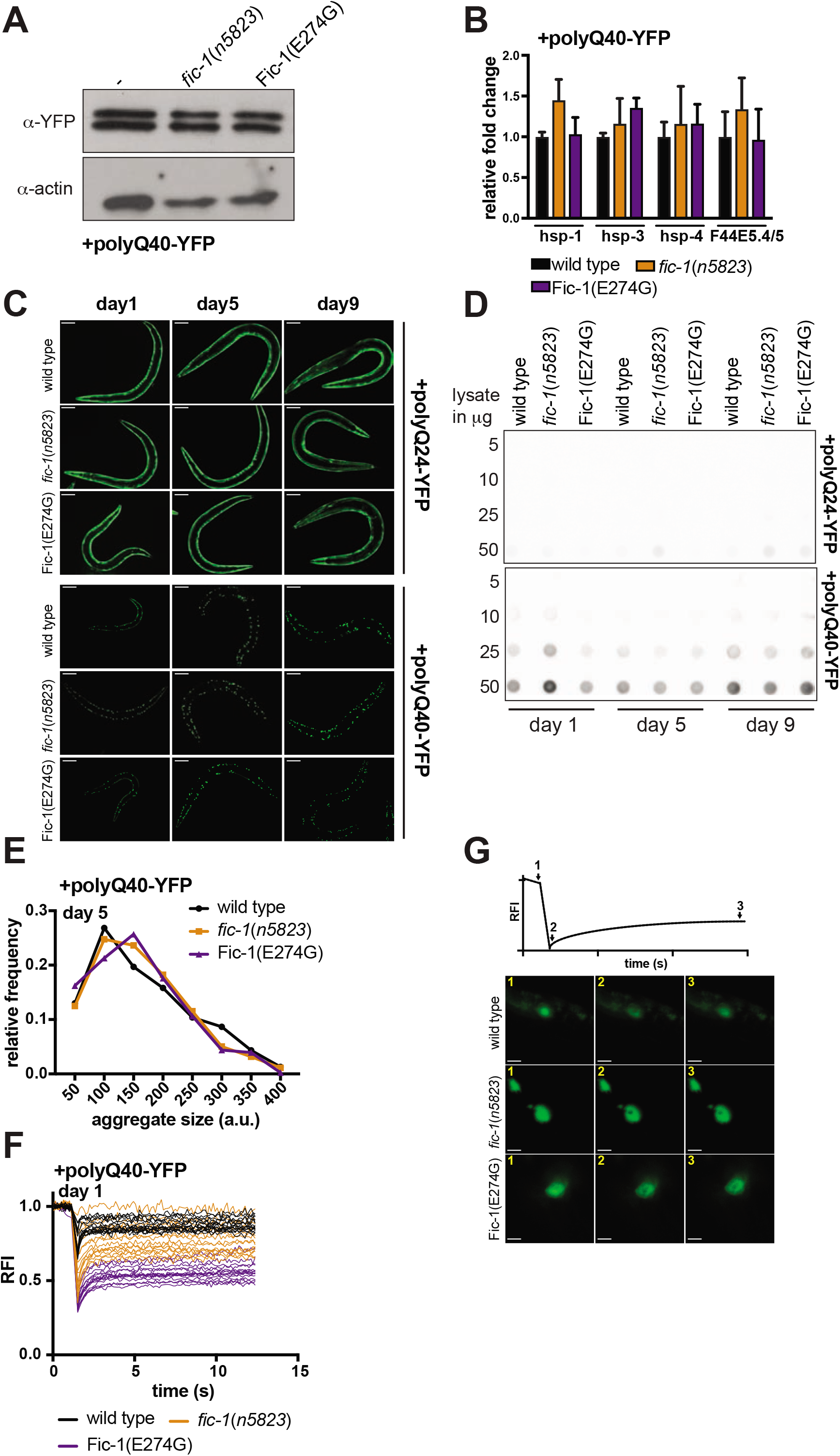
AMPylation controls aggregation of polyQ40 repeat containing proteins in *C. elegans*. (A) Immunoblot showing polyQ40-YFP abundance in indicated *C. elegans* total lysates. Total protein concentrations were determined by a Micro BCA assay and equal amounts of protein were loaded in each lane. Actin serves as additional loading control. (B) qPCR analysis of hsp-1, hsp-3, hsp-4 and F44E5.4/5 transcript abundance in wild type, *fic-1(n5823)* and Fic-1(E274G). Error bars represent SD. (C) representative micrographs showing polyQ24-YFP (panels 1-3) and polyQ40-YFP aggregation (panels 4-6) of wild type, *fic-1(n5823)* and Fic-1(E274G) at day 1, day 5 and day 9 of adulthood. Scale bar equals 100 μm (D) filter trap assay of polyQ24-YFP and polyQ40-YFP aggregates. Indicated amounts of *C. elegans* lysates were adsorbed onto cellulose acetate membranes and polQ-YFP aggregates were detected by immunoblotting. (E) distribution profile of polyQ40-YFP aggregate sizes on day 5 of adulthood. Bin size is 50 a.u. (F) polyQ40-YFP aggregate mobility on day 1 of adulthood as measured by fluorescence after photobleaching (FRAP). Y-axis depicts relative fluorescence intensity (RFΊ). (G) Schematic overview and representative micrographs of FRAP testing for day 1 adults as quantified in (F). Scale bar equals 3 μm.

**Figure S4:**
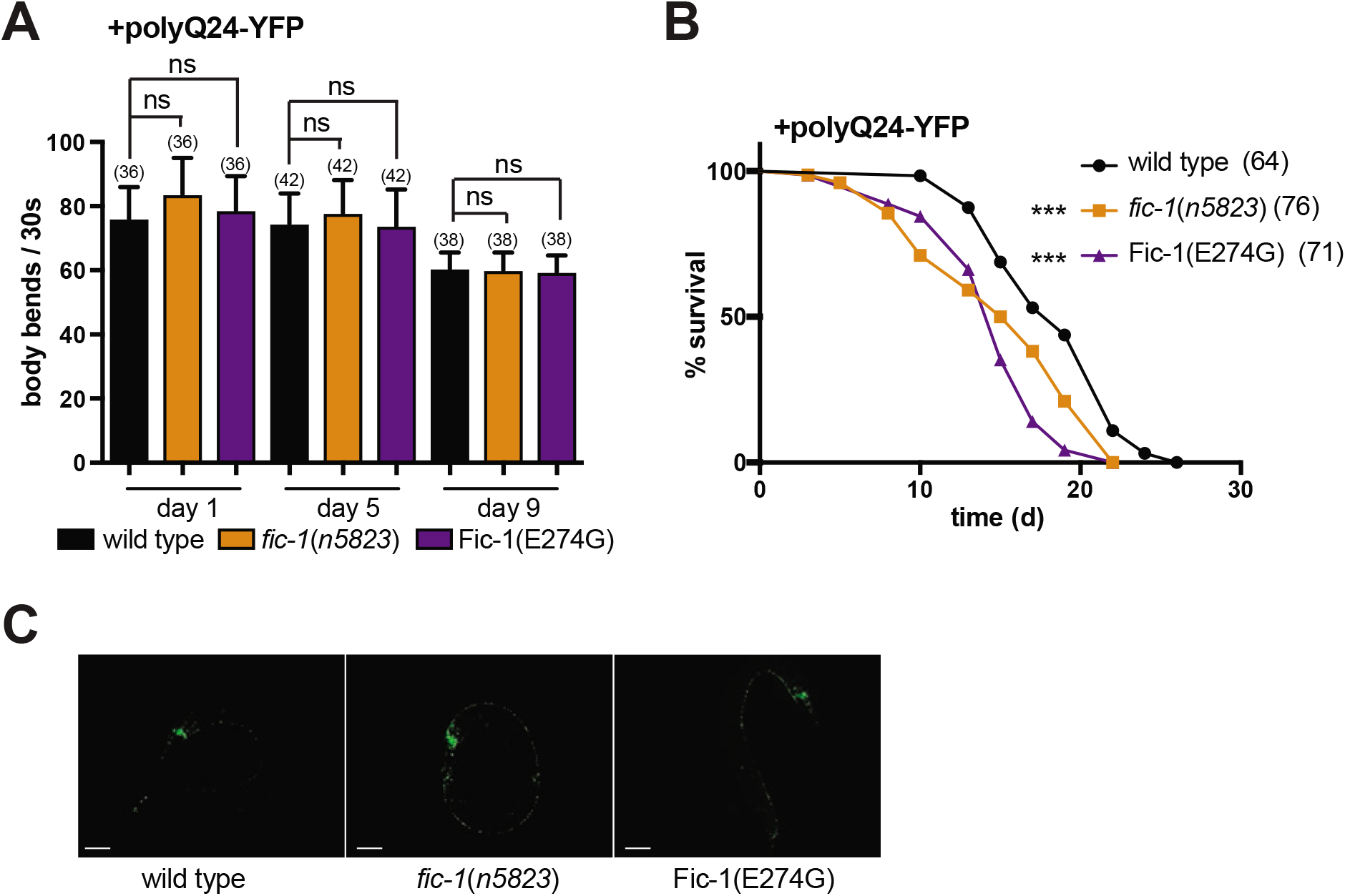
AMPylation affects polyQ-YFP toxicity in *C. elegans*. (A) Motility assay of polyQ24-YFP-expressing wild type, *fic-1(n5823)* and Fic-1(E274G) worms at the indicated time points. (B) lifespan analysis of polyQ24-YFP-expressing wild type, *fic-1(n5823)* and Fic-1(E274G) animals. For (A) and (B): Number of tested animals per group is shown in brackets. Error bars represent SD. Statistical significance (P-values) were calculated using the Mann-Whitney test as compared to N2 wild type control. **\*p < 0.05, **p < 0.01, ***p < 0.001, not significant (ns) p > 0.05.** (C) fluorescence microscopy of polyQ86-YFP-expressing wild type, *fic-1(n5823)* and Fic-1(E274G) worms at day 1 of adulthood, corresponding to aggregate quantification shown in Fig. 4D. Scale bar equals 100 μm.

**Figure S5:**
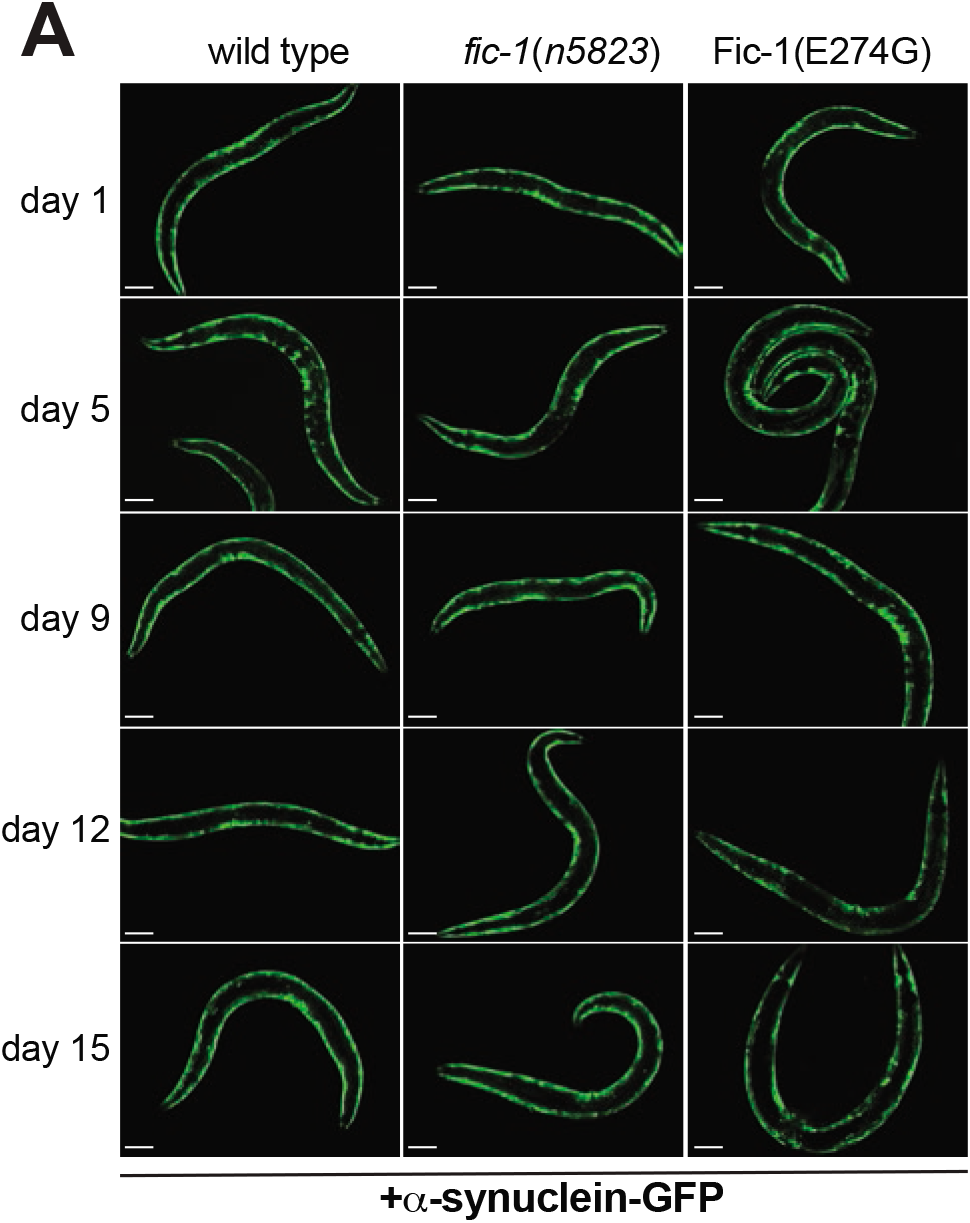
AMPylation alters α-synuclein toxicity in *C. elegans*. (A) Fluorescent micrographs showing wild type, *fic-1(n5823)* and Fic-1(E274G) worms expressing α-synuclein at the indicated age. Scale bar equals 100 μm.

**Figure S6:**
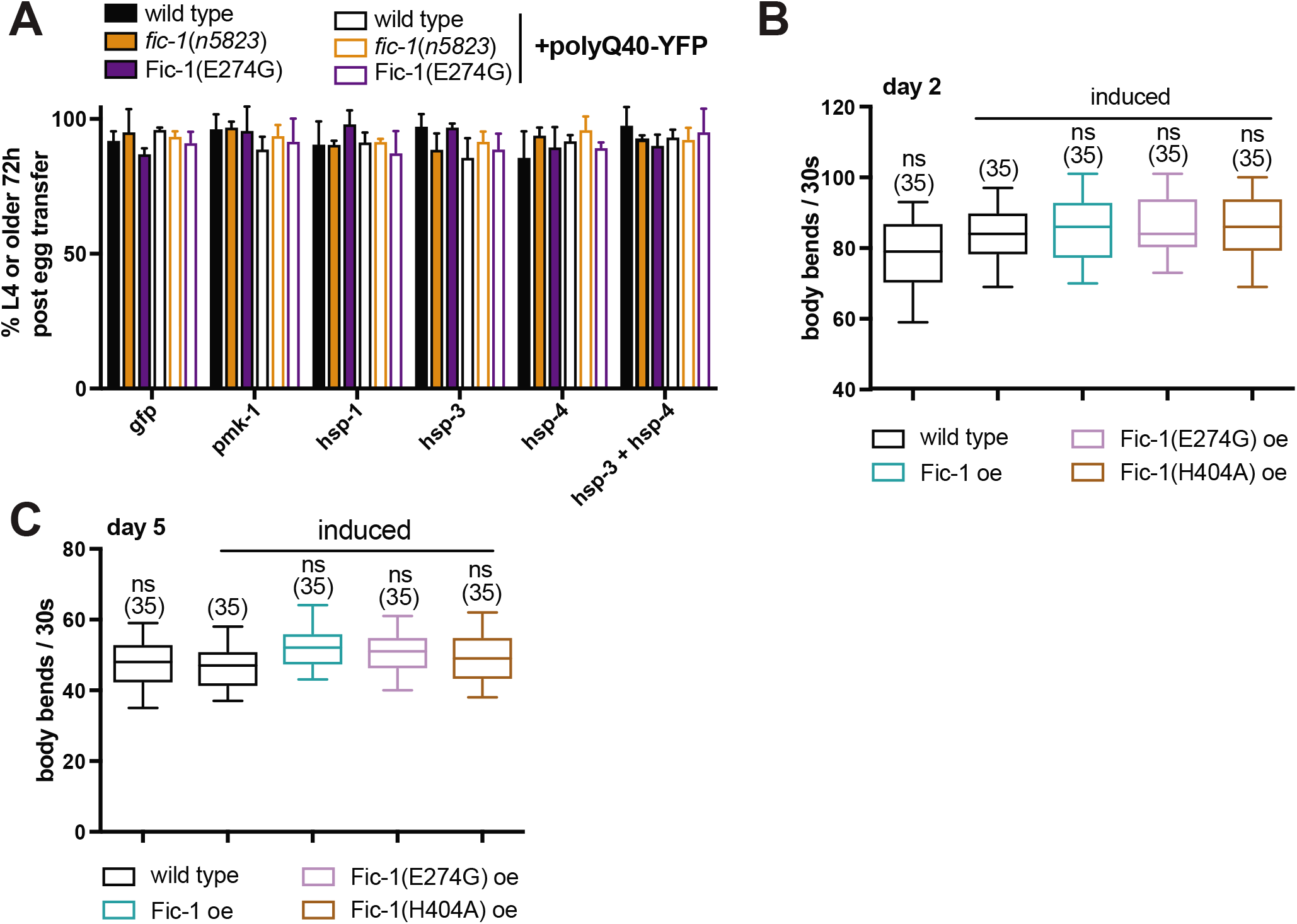
Extensive protein AMPylation interferes with larval development and heat stress tolerance in *C. elegans*. (A) Development assay depicting the proportion of wild type, *fic-1(n5823)* and Fic-1(E274G) larvae that do or do not express polyQ40-YFP and are able to reach adulthood within 72 hours at 20 °C when placed on RNAi plates as L3 larvae. (B) Immunoblot of indicated *C. elegans* total lysates. Total protein concentrations were determined by a Micro BCA assay and equal amounts of protein were loaded in each lane. HA-tagged, over-expressed proteins were detected using α-HA Ab (C-D). Motility tests of worms induced to express the indicated proteins as day 1 adults on day 2 (C) or day 5 (D). For (A) to (D): Number of tested animals per group is shown in brackets. Error bars represent SD. Statistical significance (P-values) were calculated using the Mann-Whitney test as compared to induced wild type control. **\*p < 0.05, **p < 0.01, ***p < 0. 001, not significant (ns) p > 0.05**.

